# Snipe: Highly sensitive pathogen detection from metagenomic sequencing data

**DOI:** 10.1101/2020.05.06.080580

**Authors:** Lihong Huang, Bin Hong, Wenxian Yang, Liansheng Wang, Rongshan Yu

**Affiliations:** School of Informatics, Xiamen University, China; Aginome Scientific, Xiamen, China

## Abstract

Metagenomics data provides rich information for the detection of foodborne pathogens from food and environmental samples that are mixed with complex background bacteria strains. While pathogen detection from metagenomic sequencing data has become an activity of increasing interest, shotgun sequencing of uncultured food samples typically produces data that contains reads from many different organisms, making accurate strain typing a challenging task. Particularly, as many pathogens may contain a common set of genes that are highly similar to those from normal bacteria in food samples, traditional strain-level abundance profiling approaches do not perform well at detecting pathogens of very low abundance levels. To overcome this limitation, we propose an abundance correction method based on species-specific genomic regions to achieve high sensitivity and high specificity in target pathogen detection at low abundance.

## 1 Introduction

Foodborne diseases present a notable threaten to global food safety and public health in both developed and developing countries. Every year, food contaminated with pathogens cause approximately 48 million people diseases, 128,000 hospitalizations and 3,000 deaths in the U.S.A. (www.producesafetyproject.org). The total economic burden may exceed $152 billion per year, including $39 billion due to the contamination of fresh and processed food^1^. Globally, nearly a quarter of the population is at high risk for foodborne diseases nowadays^2^.

Generally, foodborne diseases are associated with the consumption of raw or undercooked food contaminated with pathogens or their toxins. In clinical decision-making associated with foodborne diseases, the delay in diagnosis often leads to delayed treatment or inappropriate antibiotic use. In large-scale foodborne disease outbreaks, delay in detecting the specific pathogenic sources could result in an increase in the incidence and spread of disease through communities and increased mortality rates. Therefore, the ability to rapidly and accurately identify foodborne pathogens is essential to ensure food supply safety and to minimize the impact of foodborne illnesses on public health.

Conventionally, culture-based methods have been used for routine clinical diagnostic and detection of foodborne pathogens where microorganisms are cultured on agar plates, followed by standard biochemical identifications using plating techniques, bioluminescence, flow cytometry, MALDI mass spectroscopy, DEFT, and impedimetric immunoassays, etc^3, 4^. Those methods are known to be time-consuming and laborious. Culture-based methods require two to three days for initial identification and more than a week for confirmation of the pathogen species in the cultured samples^5^. In addition, since successful detection of pathogens using culture-based methods depends on the ability of the target microorganisms to grow in the culture media, sensitivity of these methods may be limited as false negative results could occur when viable but non-culturable cells are present^6^. For these reasons, traditional culture-based methods are increasingly being perceived as insufficient to meet the demands of rapid food testing and significant efforts have been invested in developing methods that are able to identify the source of foodborne diseases more accurately, more rapidly, and at lower-cost.

To speed up the pathogen detection procedure, culture-independent approaches based on quantitative polymerase chain reaction (PCR) amplification have been developed^7^. Compared to culture-based methods, PCR is faster and is able to provide real-time detection of foodborne pathogens and toxins with high sensitivity and selectivity even in the absence of a selective enrichment medium^8^. However, as PCR-based methods depend heavily on the amplification of preselected target-specific sequences, it is challenging for these methods to balance between the coverage and resolution in a single panel for closely related organisms isolated from food or environmental samples, which can be crucial to identify the right clusters for action.

With the progress of the next generation sequencing (NGS) technology and its decreasing cost, shotgun metagenomics sequencing has been rapidly adopted as the ultimate tool for microbial profiling^9,10^. By performing genome sequencing on genetic material directly collected from food or environmental samples, shotgun sequencing serves as an essential tool to gain full understanding on what pathogens are present and how they impact the food samples or environments around them, and provides the potential to deliver pathogen detection results at unprecedented speed, sensitivity and resolution.

Despite the unique advantages of high-throughput metagenomic sequencing technologies, exploring these new data for rapid species identification and strain attribution remains a big challenge due to the large volume of data produced from the sequencing machines. In addition, the taxonomic diversity within the same species and the genomic similarity among different species further complicate the problem. Numerous metagenomic algorithms have been developed to tackle the data processing challenges of metagenomic NGS data. Among them, *k*-mer based algorithms, e.g., Kraken^11^, KrakenUniq^12^, Seed-Kraken^13^, CLARK^14^ and One Codex^15^, were proposed, which compare *k*-mers of metagenomic reads with those of organisms representing a wide range of clades to achieve fast and accurate taxonomic classification. Other than *k*-mers, clade-specific marker genes-based algorithms such as MetaPhlAn2^16^ and AMPHORA2^17^ were developed, where taxonomic classification is inferred from phylogenetic distances to these marker genes. Finally, read mapping-based approaches, including MEGAN^18^, Kaiju^19^, PathoScope^20^ and Sigma^21^, infer the taxonomic composition of a sample by aligning metagenomic reads against a known database of reference genomes. To provide accurate discrimination between closely related strains of the same species in metagenomic samples, statistical models are typically involved in these approaches to assign or reassign a metagenomic read to its most likely originating genome if it could be aligned to multiple genomes.

In spite of the progress in metagenomic data processing algorithms, there is still much room for improvements toward the application of metagenomic sequencing in fast detection of target pathogenic microbes at low abundance, where false positives due to genomic similarity among strains of dominant and nondominant species remain a substantial challenge to existing algorithms. In this paper, we present Snipe (SeNsItive Pathogen dEtection), a pipeline for improving the ability of existing strain-typing tools to detect common pathogens from contaminated food samples at low abundances. Snipe is based on the concept of species-specific regions (SSRs), which are unique genomic segments that can only be found in the genomes from specific species^22^. In this pipeline, the abundance of pathogens at strain level is first estimated through a read mapping-based approach, such as Pathoscope, to map the raw metagenomic reads to a reference database that contains the genomes of a set of selected pathogens of interest. After that, the raw reads are further aligned to a panel of SSRs of the target pathogens, and the *a posteriori* probability of whether a particular strain is present in the test sample is calculated based on the observed number of the reads aligned to its respective SSR. The estimated abundance is then statistically rectified based on the *a posteriori* probability. In this way, false positives due to reads misaligned to strains absent in the samples are strongly suppressed, leading to better pathogen detection performance at low abundance levels.

We compared the performance of the Snipe pipeline when it is integrated with PathoScope2^23^ with three other strain-typing tools including PathoScope2, Kraken2^24^, and Sigma^21^ using both simulated and real-world metagenomic data with pathogen spike-in. The proposed approach out-performed other methods in terms of sensitivity to the target pathogens under a preset false discovery rate (FDR) at both species and strain levels. Importantly, with the proposed SSR-based abundance correction method, the proposed approach can detect target pathogen at a relative abundance of 0.01% or less whereas all other methods tested fail at this abundance level. An early version of this approach^25^ was used by us in participating the precisionFDA CFSAN Pathogen Detection Challenge organized by the Food and Drug Administration Center for Food Safety and Applied Nutrition (FDA CFSAN). The goal of the challenge is to identify and type low abundance *Salmonella* in naturally and *in silico* contaminated samples. Our submission took the lead in strain identification performance and received the highest scores in seven out of eight evaluation tests in this challenge (https://precision.fda.gov/challenges/2/view/results).

## 2 Materials and Methods

### Overview

Snipe is designed to enable highly sensitive detection of a predefined set of target pathogens from shotgun metagenomic samples (Fig. 3). To achieve this goal, the Snipe pipeline starts with a traditional strain profiling tool such as PathoScope2 to obtain a preliminary abundance estimation of the target pathogens at both species and strain levels. After that, the metagenomic reads are further aligned to the SSRs of different species, and the numbers of reads aligned to SSRs of each species are input to a Bayesian framework to determine the *a posteriori* probabilities of the presence of target pathogen species in the metagenomic sample. Finally, the preliminary abundance estimations are rectified based on the *a posteriori* probabilities (Methods). Through this process, erroneous high abundance estimations of nonexisting species in the original sample due to misassigned metagenomic reads will be suppressed, leading to better pathogen detection results.

### Genome database construction

We created a reference database containing 2,951 complete genomes of ten selected target pathogen species, which were downloaded from NCBI RefSeq^26^ (retrieved in February 2020; total fasta file size 12 GB). The numbers of genomes for these ten species downloaded from NCBI database were 1028, 836, 534, 201, 174, 63, 55, 29, 18 and 13, respectively (Supplementary Fig. S1). This reference database was used in Snipe, PathoScope2, Sigma, and Kraken2 in our evaluation. The Snipe’s ‘bowtie2-build’ module, PathoScope2’s ‘PathoMap’ module, the Kraken2’s ‘kraken2-build’ module, and Sigma’s ‘sigma-index-genomes’ module were used for indexing the reference genomes for these four methods, respectively, all using default parameters.

Furthermore, as PathoScope2 supports the usage of a filter database to remove reads from uninterested strains in the strain profiling process, we constructed a filter database for PathoScope2, which included approximately 13,481 bacterial genomes that were not from the 10 target species obtained from NCBI RefSeq (retrieved in February 2020; total fasta file size 51 GB).

### Generate species-specific genomic regions

The SSRs were identified for each target pathogen species using the following filtering process^22^. Briefly, for each target species, the completed genomes were fragmented into 1000 bp segments using Panseq^27^, which were subsequently clustered using cd-hit v.4.6^28^ to remove potential duplicates and paralogs using a 90% sequence identity threshold. For genomic segments that remained after deduplication, we used the online NCBI BLAST^29^ tool to check if it is species-specific or if there exist similar sequences from other species.

We first used `megablast`^30^ with “highly similar sequences” optimization, and searched the genomic segment with bacteria (taxid 2) but excluded the target species itself. We then discarded genomic segments which have no less than 80% identity to bacteria genomes excluding the target species. The second round of filtering was performed using the NCBI `blastn` tool with “some-what similar sequences” optimization. The expected threshold was set to 0.001 and the word size was set to eleven to tradeoff between query sensitivity and speed. Again, the remaining genomic segments were searched against bacteria while excluded the target species itself. Genomic segments that have no less than 80% identity to bacteria genomes excluding the target species were discarded. The third round is similar to the second one, except that we only excluded the sub-species (if subspecies exist) of the target pathogen when using `blastn`. This further enhances the specificity of the selected regions at subspecies level. For each species, the identified SSRs were used as contigs to construct a pseudo SSR genome which was further used as the reference genome for SSR read identification in Snipe.

### Abundance estimation with SSR-based rectification

For abundance rectification, the metagenomic reads are first aligned to the SSRs using Bowtie2 (v2.3.4.3) with default settings. Note that end-to-end alignment is performed by default in Bowtie2 (`--ma 0`). The sum of the numbers of reads aligned to each SSR with editing distance less than or equal to 2 is recorded as the total number of SSR reads. To increase the specificity of the algorithm, only reads with read length larger than or equal to 50 basepairs (bp) are counted.

We followed the Bayesian inference to calculate the *a posteriori* probability of alternative to null hypothesis (*H*_1_) that the target pathogen is present in the metagenomic sample based on the total number of observed reads aligned to SSRs. Formally, the *a posteriori* probability of *H*_1_ can be calculated as:

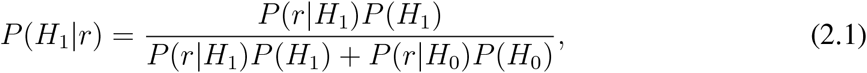

where *r* is the number of reads aligned to SSRs, *P* (*r*|*H*_1_) denotes the probability of observed *r* SSRs reads given *H*_1_. *H*_0_ is the null hypothesis that the target pathogen is not present in the sample, and *P* (*r*|*H*_0_) is thus the probability that all the reads that mapped to SSRs are misalignments due to mistakes such as sequencing error. *P* (*H*_0_) and *P* (*H*_1_) denote the *a priori* probabilities of both hypotheses.

If we further assume that each read from the metagenomic sample is an independent trial with probabilities *p*_0_ and *p*_1_ to be aligned to SSRs under hypothesis *H*_0_ and *H*_1_ respectively, the conditional probabilities *P* (*r*|*H*_0_) and *P* (*r*|*H*_1_) are binominal with parameters *p*_0_ and *p*_1_. In this case we have:

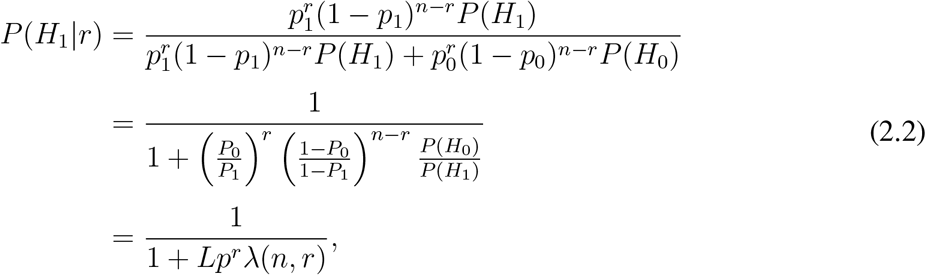

where *n* is the total number of reads from the sample, *L* ≜ *P* (*H*_0_)*/P* (*H*_1_) is the likelihood ratio of the *a priori* probabilities *P* (*H*_0_) and *P* (*H*_1_), *p* ≜ *p*_0_*/p*_1_, and

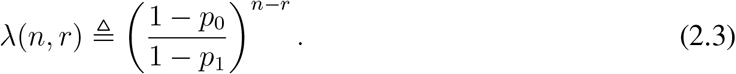

After that, the relative abundance estimations *α*_*i*_ of strain *i* is rectified with the *a posteriori* probability *P* (*H*_1_|*r*) of the target species of strain *i*, resulting in the SSRs adjusted abundance estimation 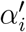 as:

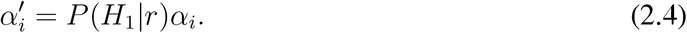

As *α*_*i*_ = 0 under the null hypothesis, 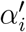 is thus the expected abundance after adjusted for additional observation of reads aligned to SSRs. In our experiment, we selected *p*_0_ = 5*e* − 10 and *p*_1_ = 2.2*e* − 5, which gave reasonable performance in our tests.

### Evaluation datasets

To create the *in silico* simulated metagenomics datasets for evaluation, we first identified a set of highly confusing microbials that were closely similar to but were not from the target pathogens. To this end, we randomly selected a reference genome from each target pathogen species, and then generated a set of 0.1 million paired-end reads of 150 bp from it using wgsim^31^ with 0.1% of mutation rate and 15% of indel fraction. The simulated reads were mapped to a subset of the NCBI’s nucleotide database (ftp://ftp.ncbi.nih.gov/blast/db/FASTA/nt.gz) using Kallisto^32^ to identify microbials that are highly similar to the selected pathogen. The top 57 most similar microbials to different target pathogen species were then combined to form a set of highly confusing background microbials. We then randomly selected a collection of ten strains from this combined set as the background strains. Similarly, we randomly selected a collection of ten strains, one from each target pathogen species, as the foreground strains.

Once the foreground and background strains were selected, we generated paired-end raw reads of 150 bp from their reference genomes using wgsim with 0.1% of mutation rate and 15% of indel fraction, and mixed the generated reads according to the desired spike-in abundance levels (0.01% - 10%) and sequencing depths (15 Mb to 1,500 Mb) to create the simulated metagenomic data. Here, the spike-in abundance level is defined as the ratio of the number of reads from the target pathogen to the total number of reads. Note that all the ten background strains were used in generating the metagenomic samples with equal abundance percentages. The strain and species composition in each *in silico* mixed metagenomic data was shown in Supplementary Fig. S11.

Finally, the metagenomic raw read data used in the precisionFDA CFSAN Pathogen Detection Challenge were downloaded from (https://precision.fda.gov/challenges/2/view). The quality of the downloaded data was assessed using fastp (v.0.19.4; http://opengene.org/fastp/fastp), and the reads were trimmed with fastp options (`-f 15 -F 15`) and (`--cut by quality 3`) to remove low quality nucleotide bases at both ends of the reads.

### Comparison to existing strain-typing tools

We compared the Snipe pipeline to PathoScope2^23^, Kraken2^24^, and Sigma^21^. PathoScope2 is a complete bioinformatics framework for quantifying the proportions of reads from individual microbial strains present in metagenomic sequencing data from environmental or clinical samples. A penalized statistical mixture model reassigns all ambiguous reads to the most probable source genome in the library. Sigma uses a short-read alignment algorithm, Bowtie2^33^, to align all metagenomic reads to every reference genome. Kraken2, a newer version of Kraken, is a taxonomic classification system that relies on exact matches of *k*-mers to the lowest common ancestor (LCA) of all genomes to achieve high accuracy and fast classification speeds. Default settings as suggested by the respective user manuals were applied when using these tools in our experiment. Abundance estimates for PathoScope2, Kraken2 and Sigma were computed from the proportion of individual reads to the total number of reads in the sample. For Snipe, the initial strain abundances were estimated using PathoScope2 with the same settings, which were further rectified based on SSR alignment results as described previously to obtain the final abundance estimation. The maximum abundances of all the strains from a particular species was taken as the estimated abundance of that species. The same reference database as described previously was used in all tools in the experiment.

We compared the performance of different tools at both strain and species levels. Here, “strain-level” accuracy is technically defined as correct pathogen identification at genome resolution. A particular pathogen detection is counted as correct at strain-level resolution if and only if the estimated abundance of its correct genome-of-origin, which is known *a priori* for simulated data or provided by data owners for external validation data, is above the detection cutoff threshold. Similarly, “species-level” accuracy is defined as correct pathogen identification at species node. A particular pathogen detection is counted as correct at species-level resolution if and only if the estimated abundances of one or more genomes attached to the same species node in the NCBI taxonomy as the correct genome-of-origin are above the detection cutoff threshold.

## 3 Result

### SSR of common foodborne pathogens

We examined the pan-genomes of ten common food-borne pathogen species^34^ (Supplementary Fig. S1) and identified genomic regions of 1000 bp that are unique to each specific species as its SSRs (Methods). The number of SSRs of different species vary from the highest 2377 SSRs in *Clostridium perfringens* to the lowest 98 SSRs in *Escherichia coli* (Fig. 1a). Considering the SSR coverage ratio, which refers to the ratio between the total length of SSRs in a species and its median reference genome length, *Escherichia coli* has the lowest SSR coverage ratio with only 1.91% of its whole genome being unique and *Clostridium perfringens* has the highest coverage ratio (69.65%). The coverage ratios of SSRs range from 3.1% to 27.76% for the other species.

**Fig. 1.**
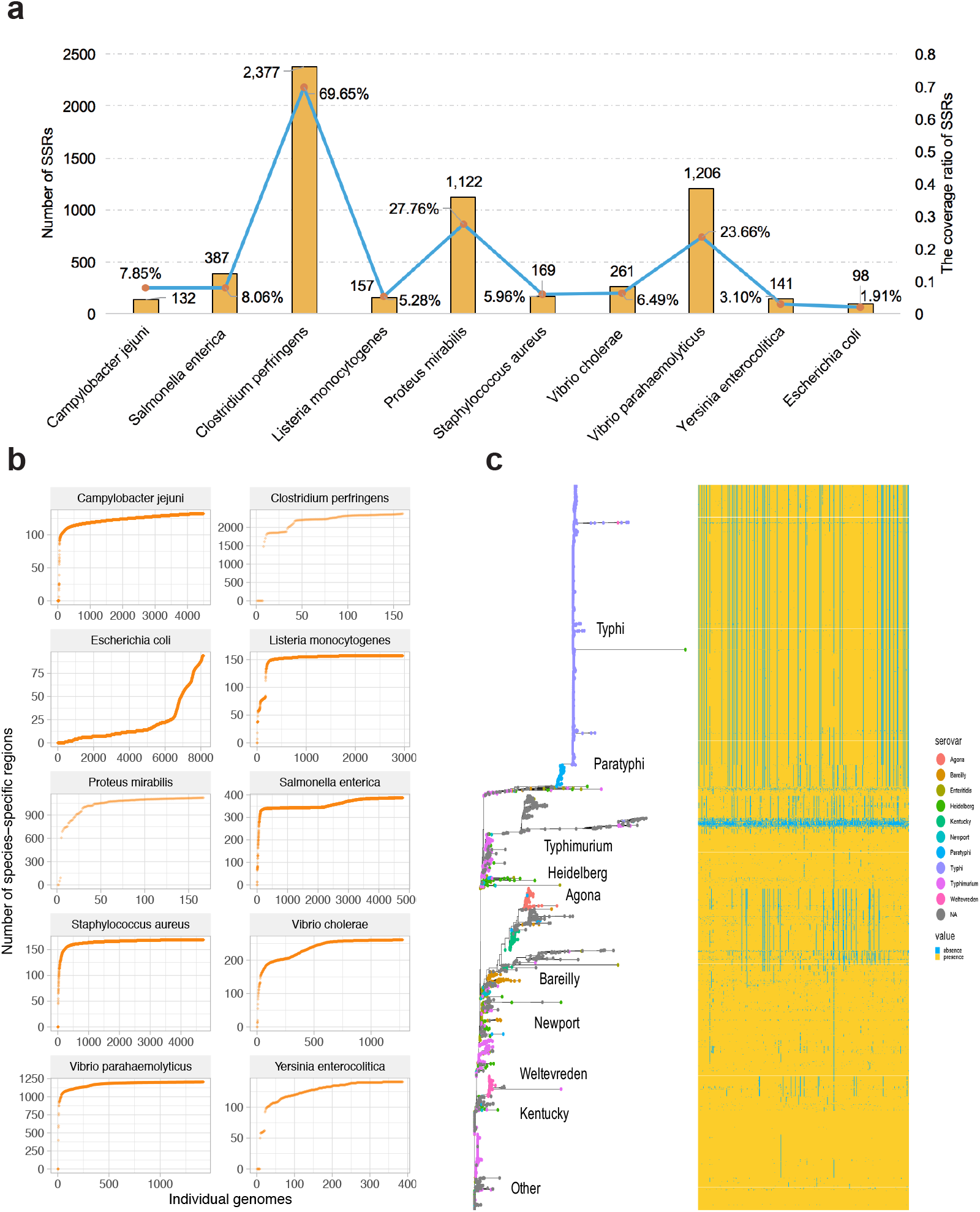
SSR of target pathogen species. **a**, The number of SSRs identified in ten species. Blue line describes the coverage ratio of SSRs with respect to median total genome length. Yellow bar shows the number of SSRs for each species. **b**, Comparison and analysis of the number of SSRs for ten common foodborne pathogen species. Each dot represents a single genome, sorted by the number of SSRs identified from that genome. **c**, The phylogeny based on the presence/absence of 1000 bp SSRs among the entire pan-genome of *Salmonella enterica*. The matrix on the right of the phylogeny shows the SSRs of *Salmonella enterica* of individual strains. Yellow represents the presence of a region, while blue represents the absence of a region.

To further understand the distribution of SSRs within each species, we analyzed the distribution of SSRs among different strains of that species. Fig. 1b shows the rather steep distributions of SSRs among different strain genomes from the same pathogen species. For most species, a significant portion of genomes contain 90% or more SSRs of the total SSRs of that species, and only a small number of genomes contain significantly less SSRs than other genomes. The only exception is *Escherichia coli*, where only a small number of genomes contain significantly more SSRs compared to the remaining genomes. We further analyzed the phylogeny of each species based on the presence or absence of SSRs in individual strain genomes. Fig. 1c shows the distribution of SSRs in 4939 genomes from different serovar types of *Salmonella enterica* as an example. The genomes for most of the serovar types in *Salmonella enterica* contain a significant number of SSRs. Strains containing small numbers of SSRs can be found mostly in the cluster located in the middle part of the figure. The corresponding serotypes of those strains are either not available or unknown, suggesting the possibilities of database labeling errors or incomplete genomes for those genomes. Similar results can be observed in the distribution of SSRs in other species as well (Supplementary Fig. S2 to Fig. S10).

### Pathogen detection at extremely low abundance levels with SSR-based abundance rectification

First, we evaluated the ability of existing strain profiling tools to identify target pathogens at low abundance from metagenomics data. To this end, we used ten simulated raw metagenomics sequencing data spiked with a different known foodborne pathogen at relative abundance of 0.01% (Methods). We then studied the number of reads being correctly attributed to the spike-in pathogens at strain and species levels for three existing tools, namely, PathoScope2, Sigma, and Kraken2. A database of 2,951 complete genomes, including genomes of the spike-in pathogens, was used as the reference strain database in all tools (Methods).

All the three tools were able to assign reads to the correct spike-in pathogen at species level at this low abundance (Fig. 2). PathoScope2 achieved the highest sensitivity and assigned more reads compared to Kraken2 and Sigma. At the strain level, PathoScope2 assigned reads to the correct spike-in pathogen strain in eight out of the ten samples, while Sigma only assigned reads to the correct spike-in pathogen strain in two samples. However, for all the three tools, a significant amount of reads were mistakenly assigned to decoy genomes that were not present in the metagenomic data (Fig. 2). In fact, in most cases, the highest read count from decoy genomes was even significantly higher than that of the correct spike-in pathogen. Therefore, it is very challenging for those tools to correctly identify the spike-in pathogens at low abundance levels due to high amount of false positive read assignments.

**Fig. 2.**
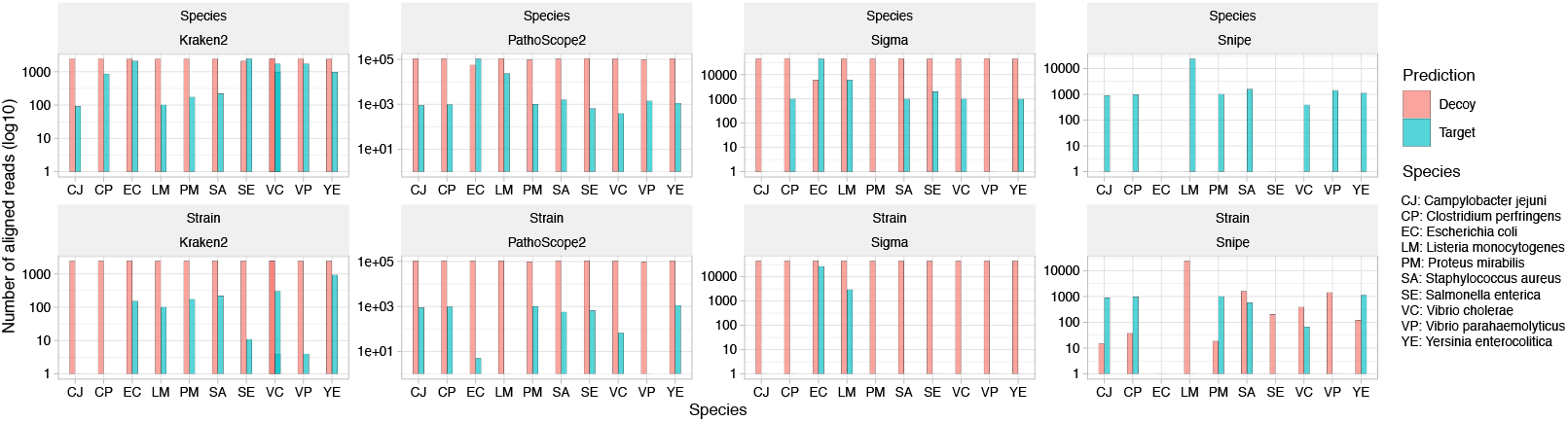
Read assignment of low abundance spike-in pathogen. Number of raw metagenomic reads assigned to the reference genomes of the correct spike-in pathogens by PathoScope2, Sigma, Kraken2 and Snipe on simulated metagenomic data with pathogen spike-in at relative abundance level of 0.01% (sequencing depth = 1,500 Mb). Blue bars indicate the number of reads assigned to the correct genome of spike-in strain (top) or species (bottom) and green bars indicate the numbers of reads assigned to strain or species not present in the simulated data (decoy strain/species) with the highest numbers of assigned reads. Number of assigned reads from Snipe were rectified by the *a posteriori* probabilities of the presence of target species calculated based on the number of SSR-aligned reads.

**Fig. 3.**
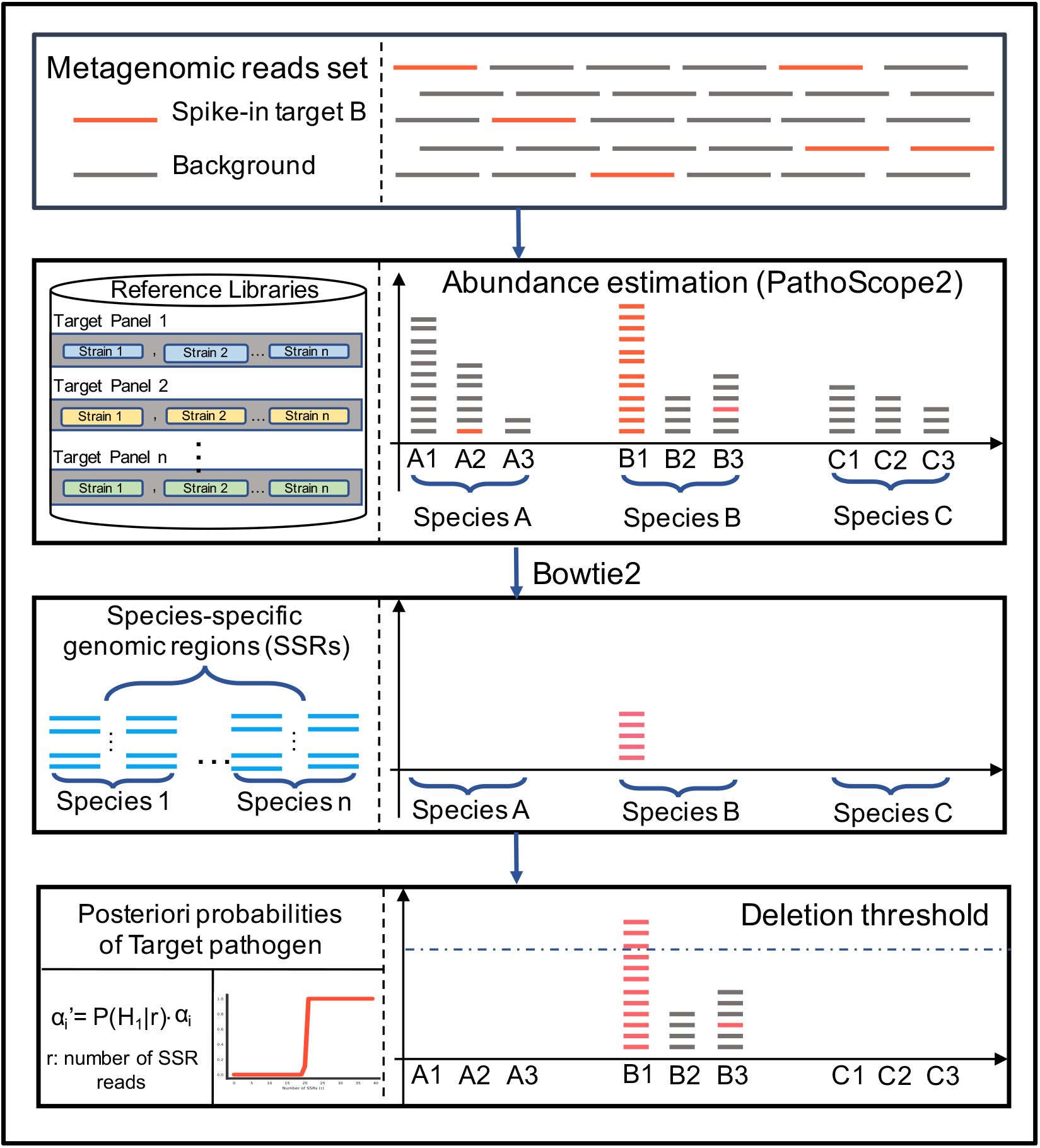
Snipe pipeline. Snipe for ultra sensitive pathogen detection. First, raw metagenomic reads are input to a strain-typing tool such as PathoScope2 to obtain a preliminary abundance estimation of target pathogens in the reference database. Then, the raw metagenomic reads are aligned to SSRs by Bowtie2 to obtain the number of SSR-aligned reads of different species, which are further used to calculate the *a posteriori* probabilities of the presence of different target pathogen species in the metagenomic sample. Finally, the preliminary abundance estimations from PathoScope2 were rectified by the *a posteriori* probabilities to obtained the final abundance estimation of target pathogens at both species and strain levels.

We further performed *a posteriori* probabilities assisted abundance rectification (Methods) on results from Pathoscope2 (the Snipe pipeline). It can be seen that the rectification step successfully reduced the number of misassigned reads while kept the number of correctly assigned reads intact in most cases (Fig. 2), indicating a better chance to correctly identify the target pathogens for strain-typing tools with the proposed abundance rectification method.

### Performance evaluation on simulated metagenomic datasets

To further evaluate the pathogen detection performance of Snipe, we created a set of 200 simulated metagenomic samples with spike-in’s from ten common foodborne pathogen species at different relative abundance levels and sequencing depths (Methods). We then profiled these samples with PathoScope2, Kraken2, Sigma and Snipe, and evaluated their sensitivity in terms of correctly identifying the spike-in pathogens at both species and strain levels. A pathogen was considered to be positively identified if its estimated abundance is above the cutoff threshold, which was determined individually for each method based on the estimated abundances of decoy species/strains on the simulated dataset such that the FDR on simulated dataset is lower than a preset threshold.

We compared the performance of Snipe versus other microbial profiling tools in identifying the spike-in target pathogens at different relative spike-in abundance, ranging from 0.01% to 10% (Fig. 4a). The result demonstrated superior sensitivity performance of Snipe at both species and strain levels. Particularly, at the lowest abundance level of 0.01%, all the other methods failed to identify almost all the spike-in pathogens, while Snipe was still able to correctly identify all the spike-in pathogens at species level. In addition, for 60% of the simulated samples, Snipe correctly identified the strains of the spike-in samples at the sequencing depth of 1,500 Mb. Note that as the number of reads captured by SSRs is proportional to the total number of reads of the spike-in pathogens, the sensitivity performance of Snipe is dependent on the sequencing depth of the metagenomic samples. As we reduced the sequencing depth to 15 Mb, the sensitivity of Snipe reduced to 50% and 10% at species and strain levels, respectively. Furthermore, we noticed that the sensitivity performance of Snipe saturated at a sequencing depth of 150 Mb. Other strain profiling methods, such as PathoScope2, also benefit from increasing sequencing depth, but to a lesser extent. At the highest spike-in abundance level of 10%, all the methods except Kraken2 correctly identified all the spike-in pathogens at both species and strain levels, even at the lowest sequencing depth of 15 Mb. We further broke down the pathogen identification results from different tools according to different species (Fig. 4b-c). The most challenging species for Snipe in strain identification were *Listeria monocytogenes* and *Escherichia coli*, where correct strain identification was achieved only at spike-in abundance of 1% or higher. For other species, Snipe showed excellent performance even at the lowest spike-in abundance (0.01%).

**Fig. 4.**
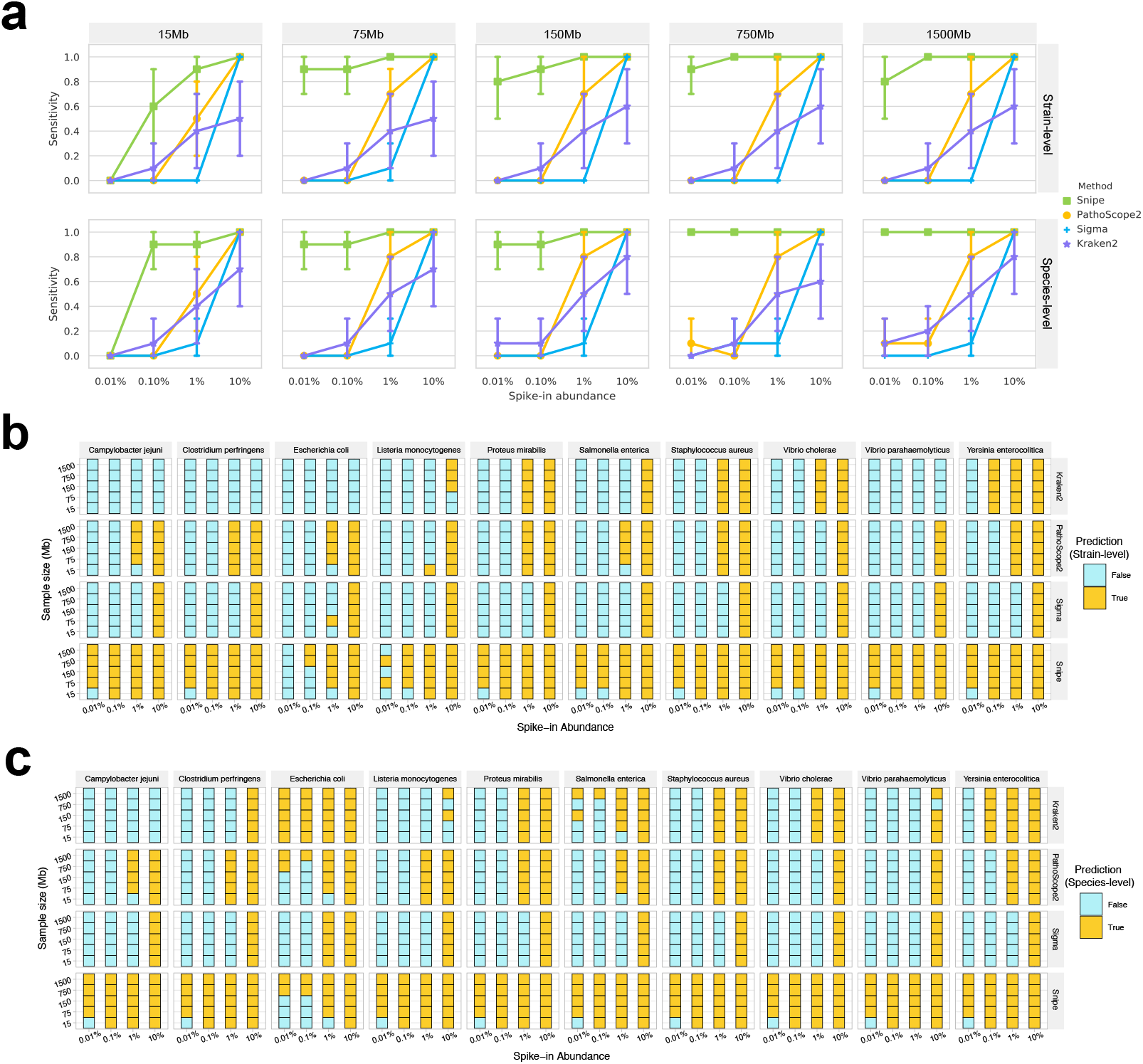
Low abundance pathogen detection enabled by SSR-based rectification. **a**, Pathogen detection sensitivity of different strain profiling tools for four spike-in abundance from 0.01% to 10% at different sequencing depth from 15 Mb to 1,500 Mb. Center line is the averaged sensitivity over ten target pathogen species under test, and lower and upper whiskers indicate 95% confidence interval. **b-c**, Comparison of pathogen identification results of Snipe, PathoScope2, Sigma, and Kraken2 from 200 simulated data at 5 different abundance levels from 10% to 0.01% at species (**b**) and strain (**c**) levels. Yellow square means that the method successfully identifies the target pathogen from the simulated sample data, while blue square means otherwise.

As Snipe depends on reads from SSRs for target identification, we asked whether the coverage ratio of SSRs with respect to the whole genome size of a species affects the pathogen identification capability of Snipe at low abundance. To this end, we plotted the sensitivity of Snipe with respect to the coverage ratio of SSRs of different species (Fig. 5), which shows strong associations between them when both the spike-in abundance and the total number of reads are low. In such case, *Clostridium perfringens*, which has the highest coverage ratio of SSRs, also achieves the highest sensitivity among all species. On the other hand, *Escherichia coli* and *Listeria monocytogenes*, which have SSRs coverage ratio of 1.91% and 5.28% respectively, are among the species that showed worst sensitivity. This result is as expected because a larger ratio indicates that the species is more distinguishable from other species, and hence easier to detect at low abundance.

**Fig. 5.**
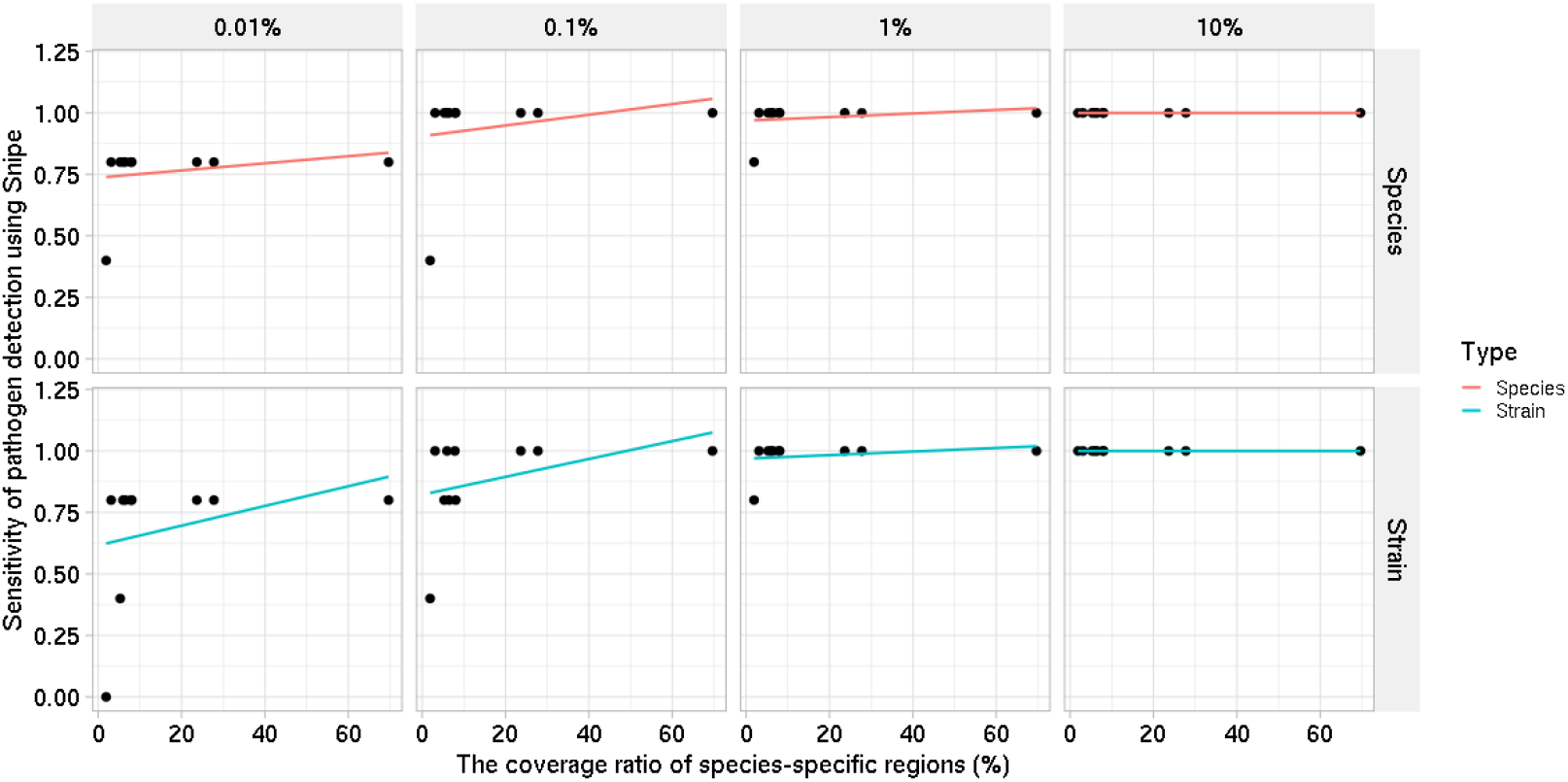
The relationship between pathogen detection sensitivity and the number of SSRs of the target pathogen. The coverage ratio (%) denotes the ratio between the total length of SSRs and the median total length of the reference genome of the target pathogen.

### Demonstration of Snipe on FDA pathogen dataset

We further assessed the performance of Snipe on 24 samples of the precisionFDA-provided dataset available from the Pathogen Challenge (https://precision.fda.gov/challenges/2/). Among the 13 positive samples, five samples were created synthetically by adding *Salmonella* reads to culture-negative samples and eight were real-life samples contaminated with *Salmonella*. The cutoff values of all tools for detection were selected based on an FDR smaller or equal 10% on our simulated set.

The identification results are shown in Fig. 6. It can be seen that Snipe successfully identified eleven positive samples at species level, and correctly identified seven positive samples at strain level. Kraken2 achieved the second-best sensitivity performance on this dataset, and correctly identified eleven out of 13 samples at species level. Unfortunately, it failed to provide any strain-level information on those positive samples. Sigma showed the worst performance and only identified eight out of 13 samples at species level. PathoScope2 also correctly identified seven positive samples at strain level. However, it missed three cultured-positive samples at species level. Interestingly, for sample C24 which has a relatively high *Salmonella enterica* abundance as indicated by its high number of SSR reads, all the four tools identified the target strain as Levine 15 instead of the groundtruth MOD1 SALC 120 strain although both strains exist in the reference database. Similarly, sample C08 was identified as FL_FLDACS-98213 by Patho-Scope2, Sigma, and Snipe, instead of its groundtruth MOD1 SALC 120 strain. Further analysis showed that the MOD1 SALC 120 and the Levine 15 strains have an average nucleotide identity^35^ (ANI) of 99.98% between their genomes, and the MOD1 SALC 120 and the FL FLDACS-98213 strains have an ANI of 98.32%. Moreover, using Bowtie2 as aligner, we found that 642,709 reads from C24 were uniquely aligned to Levine 15, while only 5,258 reads were uniquely aligned to MOD1 SALC 120. For C08, 430,752 reads were uniquely aligned to FL FLDACS-98213, and only 59 reads were uniquely aligned to MOD1 SALC 120. Therefore, it is possible that either these two samples were mislabeled, or both strains co-existed in the original samples.

**Fig. 6.**
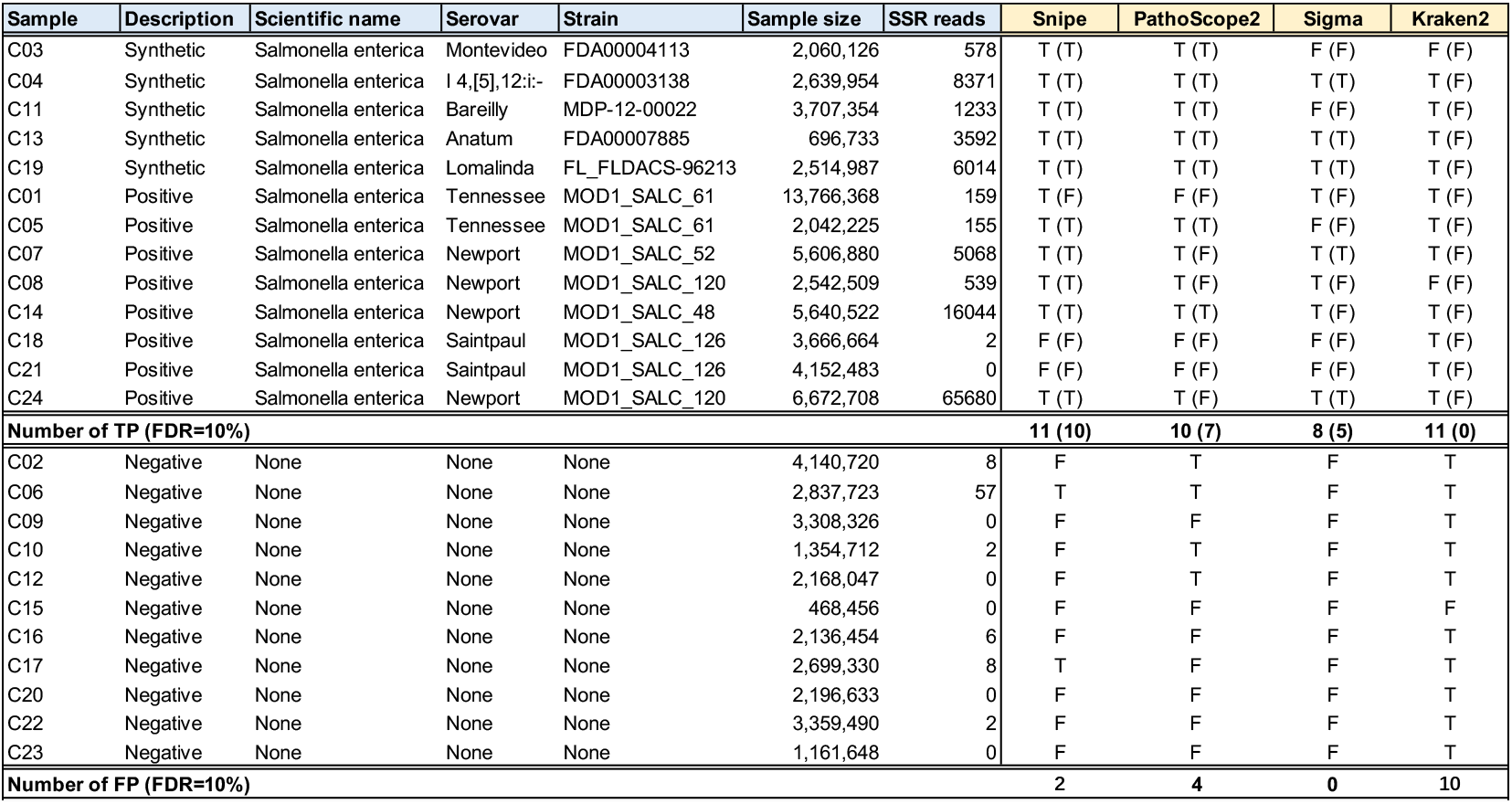
Comparison of four methods on PrecisionFDA CFSAN Pathogen Detection Challenge samples. SSR read denotes the number of reads that were aligned to SSRs of *Salmonella enterica* by Bowtie2 with editing distance smaller than or equal to three. For each method, in cell, “T” (True) represents *Salmonella enterica* was identified with estimated abundnace higher than the cutoff value determined to satisfy FDR ≤ 10% on our simulation dataset for each method, and “F” (False) represents otherwise, i.e., either that *Salmonella enterica* is not identified by the profiling tool, or the estimated abundance is below the cutoff value. Furthermore, for culture-positive or synthetic samples, “T” in parenthese indicates that the groundtruth strain as provided by FDA was identified with estimated abundance higher than the cutoff value, and “F” indicates otherwise.

In the eleven negative samples from this dataset, Snipe labeled only one sample (C06) as *Salmonella enterica* positive, which was less than PathoScope2 (four samples) and Kraken2 (ten sample). Sample C06 contains certain amount of SSR reads of *Salmonella enterica*. Manual inspection using Blast confirmed that those reads are highly specific to *Salmonella enterica*, indicating that C06 may indeed has a very low abundance of *Salmonella enterica*. Only one sample (C15) was correctly classified by all the four tools as *Salmonella enterica* negative.

### Computing time

We evaluated the run time performance of different tools using three different metagenomic samples of 0.1, 1 and 10 million reads (Fig. 7). For all methods, the experiment was implemented on an Intel® Xeon(R) workstation with 40 CPU threads and 64 GB RAM. All tools were tested with their default parameters and 32 CPU threads. For fairness, the same reference database was used in all tools. Kraken2 is the fastest among all tools due to its direct mapping from *k*-mers to the LCA of the genomes^11^. Compared to Kraken2, PathoScope2 and Sigma were much slower as substantial CPU time was used to assign reads to genomes in the reference database through read alignment and statistical inference, which were computationally intensive but necessary if one wishes to attribute each metagenomic read to its organism of origin at strain level. Compared to other methods, Snipe pipeline adds an additional overhead of aligning reads to the SSRs for abundance rectification using Bowtie2. Nevertheless, as the sizes of SSRs are significantly smaller compared to those of the full genomes in the reference database, the overhead is small.

**Fig. 7.**
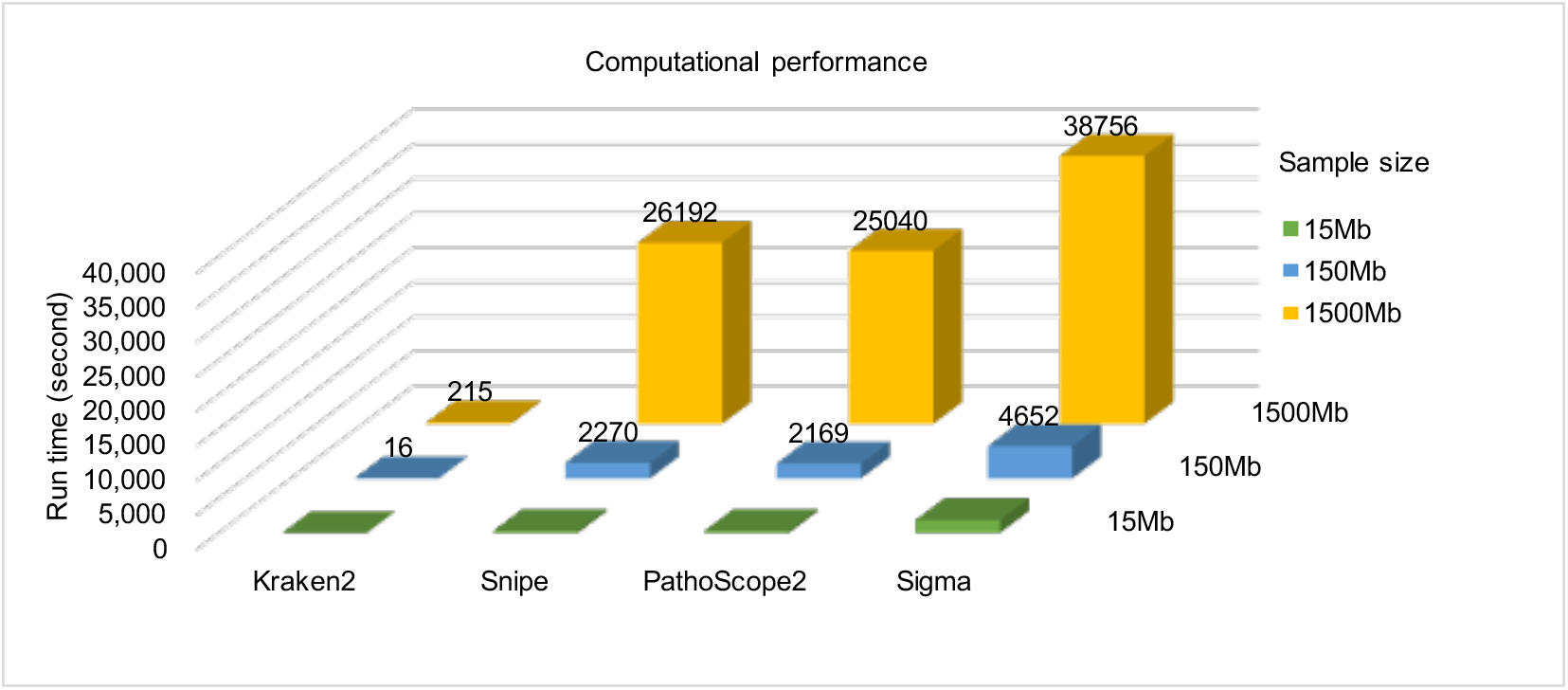
Run time performance of four tools (Bowtie2, Kraken2, PathoScope2, and Sigma). Run time performance was evaluated on the simulated metagenomic datasets at different sequencing depth of 15 Mb - 1,500 Mb. Note that the run time of Snipe is the total run time of Bowtie2 for read alignment to SSRs and PathoScope2 for initial strain abundance estimation.

## 4 Discussion

We presented a highly efficient and sensitive computational approach for detecting pathogens from metagenomics data at very low abundance levels, where the associations of metagenomic reads with target pathogens are difficult to be discovered due to reads misaligned to microbial strains that are not present in the sample. Snipe effectively suppresses a significant portion of such noise by rectifying the abundance estimations of target pathogens with their *a posteriori* probabilities inferred from the number of reads captured in genome regions that are unique to these targets, thus significantly increases the pathogen detection sensitivity without compromising the specificity performance.

Note that Snipe, to some extent, resembles the concept of other nucleic acid detection based methods, where specific nucleic acid targets are amplified or enriched by technologies such as PCR, nucleic acid sequence-based amplification (NASBA), loop-mediated isothermal amplification (LAMP) or oligonucleotide DNA microarray for time-efficient and sensitive pathogen detection at low abundance^7^. Here, instead of restricting the detection to a list of limited number of specific targets, we extensively analyzed the available pan-genomes from GenBank databases maintained by the National Center for Biotechnology Information (NCBI) to identify a comprehensive set of specific genome regions of the target pathogens, which are then used in combination with shotgun metagenomic sequencing to enable unbiased detection of a broad spectrum of pathogens at strain-level resolution and extremely low abundance levels. Thus, Snipe combines the merits of both untargeted metagenomic NGS with target-based approaches to deliver the sensitivity, resolution and coverage that are not simultaneously achievable for conventional approaches.

The proposed approach has limitations. First, the detection sensitivity of Snipe is bounded by the availability of SSR reads from the metagenomic data. As the distribution of number of SSRs of different strains from a same species can be highly unbalanced, e.g., in *E. Coli*, Snipe may be biased towards strains with large number of SSRs. However, as illustrated in our experiments, such unbalance is diminishing at high sequencing depth when a sufficient number of SSR reads from target pathogens are available in the metagenomic sample. Another limitation of the proposed approach, as other mapping-based profiling approaches, is that previously uncharacterized pathogens are difficult to detect. In addition, the specificity of Snipe may be affected by the database bias issue as some genomic regions may be wrongly identified as SSRs if they highly overlapped with those from underrepresented species in the current NCBI database. With the rapidly increasing number of reference genomes being generated each year, including those from difficult-to-grow species by using new cultivation methods, single-cell sequencing approaches or metagenomic assembly^36^, successful application of the proposed approach in both clinical and laboratory setups is guaranteed for an increasing number of pathogens with highly diversified reference genomes available.

We demonstrated the excellent computational performance of Snipe on a set of foodborne pathogens for both *in silico* mixed samples and real-life metagenomic samples. We anticipate that this approach would complement existing strain-typing tools to extend their operability on target pathogens at extremely low abundance levels with the novel concept of SSR-based abundance rectification. This approach can be extended to include other types of pathogen or other application scenarios such as infectious disease control, analysis of unpurified environmental samples for bio-forensics and etc. Extending the Snipe pipeline to support broader applications thus remains an important direction for future development.

## Data availability

Publicly available datasets were analyzed in this study. This data can be found from https://precision.fda.gov/challenges/2/view.

## Code availability

The code is available at https://github.com/xmuyulab/Snipe.

## Acknowledgements

We thank Kairui Mao for fruitful discussions and valuable comments on this paper.

