## Supplementary for "Snipe: Highly sensitive pathogen detection from metagenomic sequencing data"

### Supplementary material for “Snipe: Highly sensitive pathogen detection from metagenomic sequencing data”

L. Huang et al.

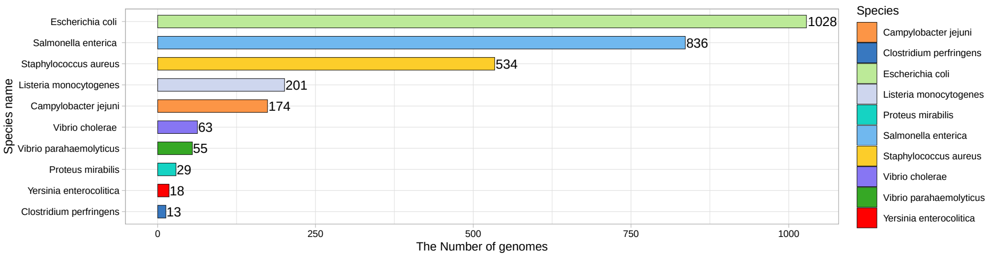


Figure S1: Numbers of complete genomes of the ten common food-borne pathogen species used in the reference database (total $n = 2,951$).

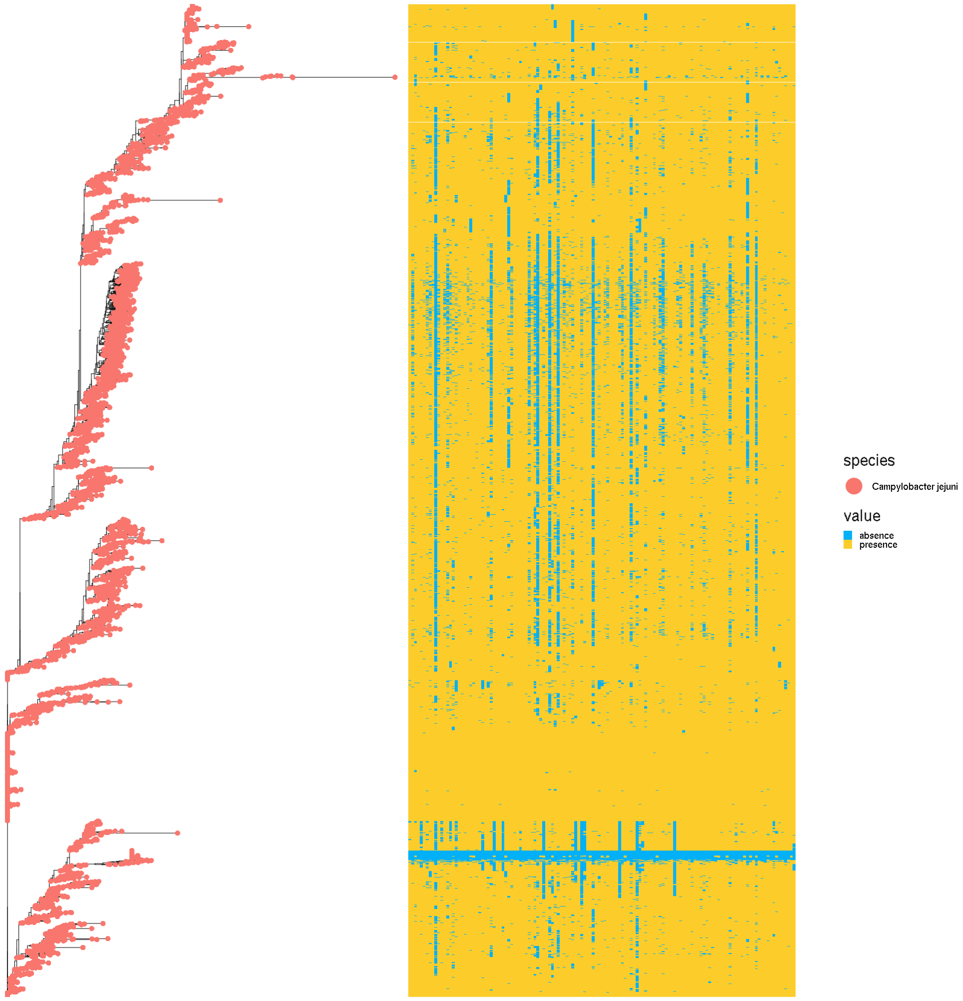


Figure S2: The phylogeny based on the presence/absence of 1000 bp SSR fragments among the entire pan-genome of *Salmonella enterica*. The matrix on the right of the phylogeny shows the SSRs of *Campylobacter jejuni* of individual strains. Color yellow represents the presence of a region, while blue represents the absence of a region.

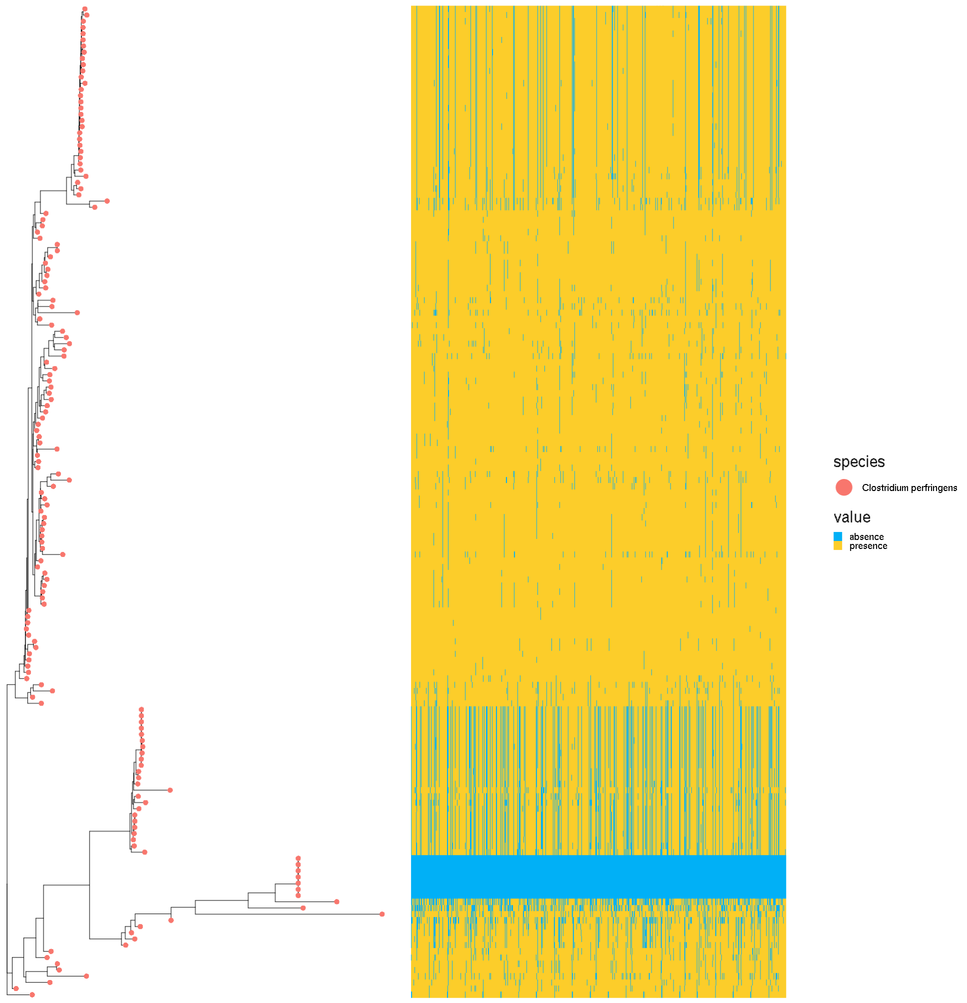


Figure S3: The phylogeny based on the presence/absence of 1000 bp SSR fragments among the entire pan-genome of *Salmonella enterica*. The matrix on the right of the phylogeny shows the SSRs of *Clostridium perfringens* of individual strains. Color yellow represents the presence of a region, while blue represents the absence of a region.

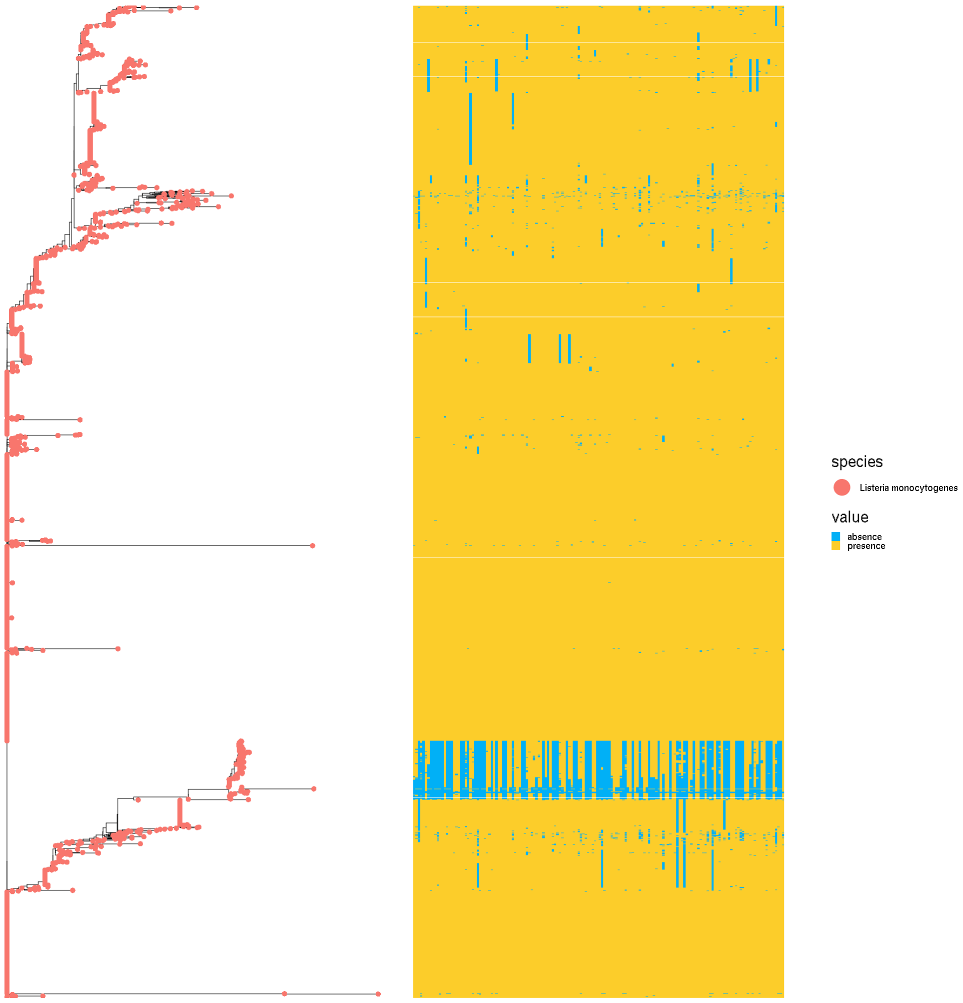


Figure S4: The phylogeny based on the presence/absence of 1000 bp SSR fragments among the entire pan-genome of *Salmonella enterica*. The matrix on the right of the phylogeny shows the SSRs of *Listeria monocytogenes* of individual strains. Color yellow represents the presence of a region, while blue represents the absence of a region.

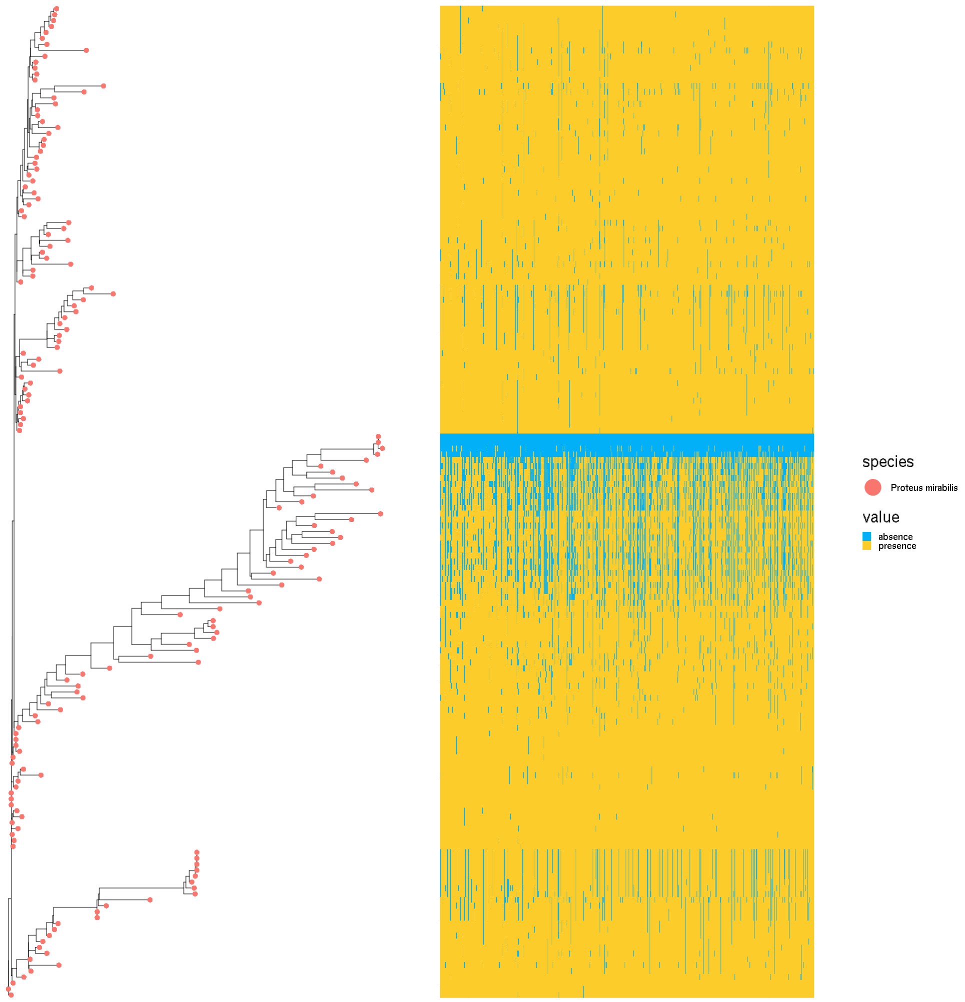


Figure S5: The phylogeny based on the presence/absence of 1000 bp SSR fragments among the entire pan-genome of *Salmonella enterica*. The matrix on the right of the phylogeny shows the SSRs of *Proteus mirabilis* of individual strains. Color yellow represents the presence of a region, while blue represents the absence of a region.

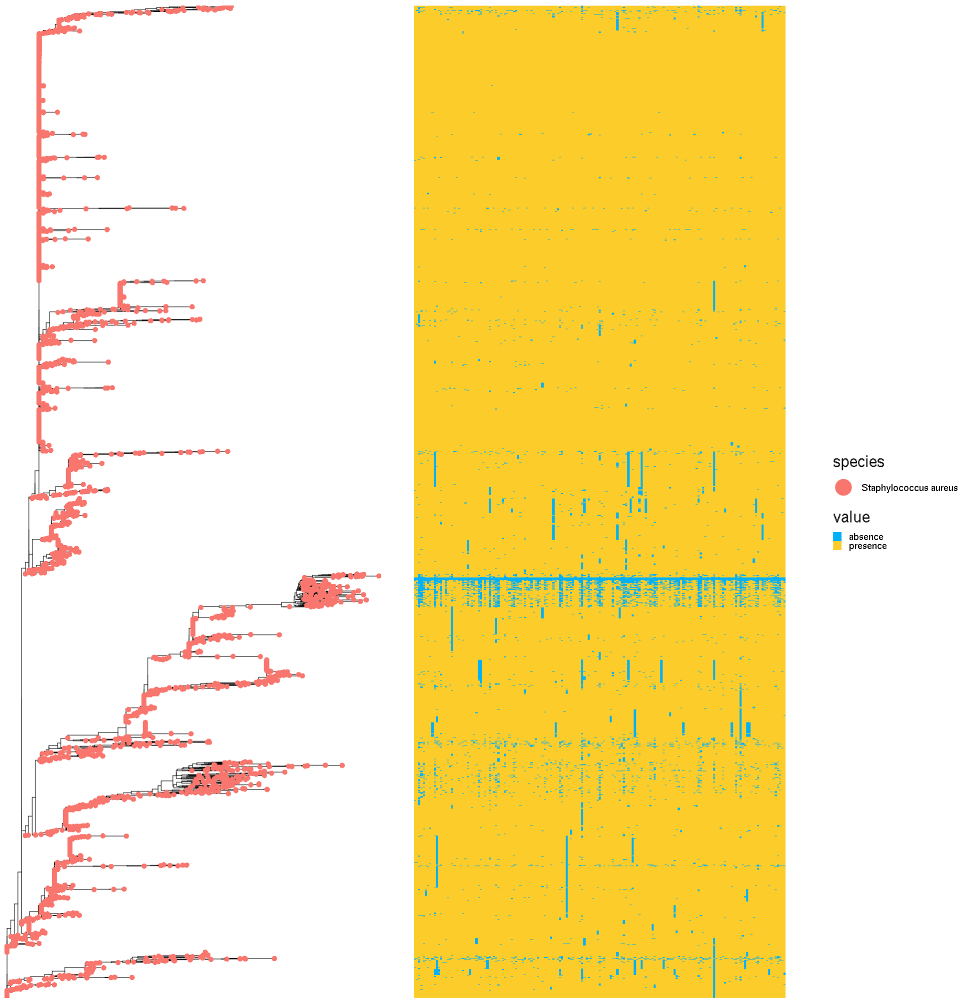


Figure S6: The phylogeny based on the presence/absence of 1000 bp SSR fragments among the entire pan-genome of *Salmonella enterica*. The matrix on the right of the phylogeny shows the SSRs of *Staphylococcus aureus* of individual strains. Color yellow represents the presence of a region, while blue represents the absence of a region.

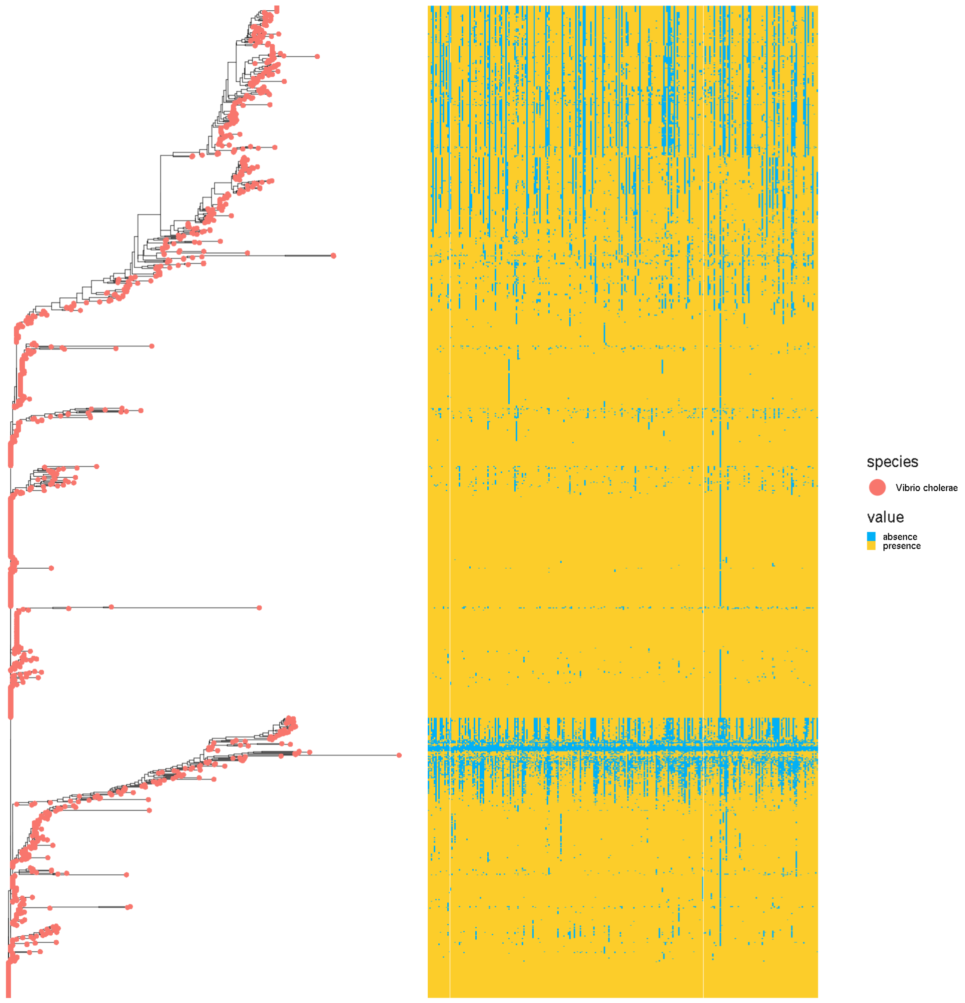


Figure S7: The phylogeny based on the presence/absence of 1000 bp SSR fragments among the entire pan-genome of *Salmonella enterica*. The matrix on the right of the phylogeny shows the SSRs of *Vibrio cholerae* of individual strains. Color yellow represents the presence of a region, while blue represents the absence of a region.

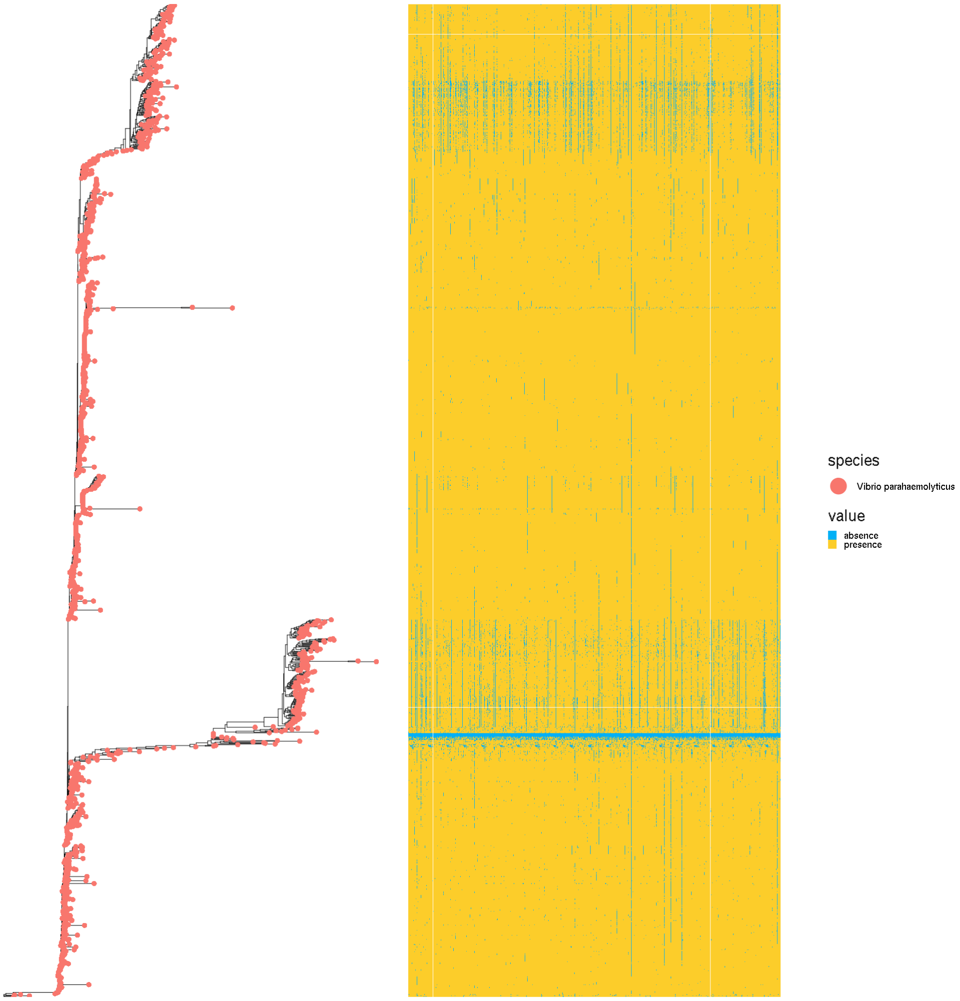


Figure S8: The phylogeny based on the presence/absence of 1000 bp SSR fragments among the entire pan-genome of *Salmonella enterica*. The matrix on the right of the phylogeny shows the SSRs of *Vibrio parahaemolyticus* of individual strains. Color yellow represents the presence of a region, while blue represents the absence of a region.

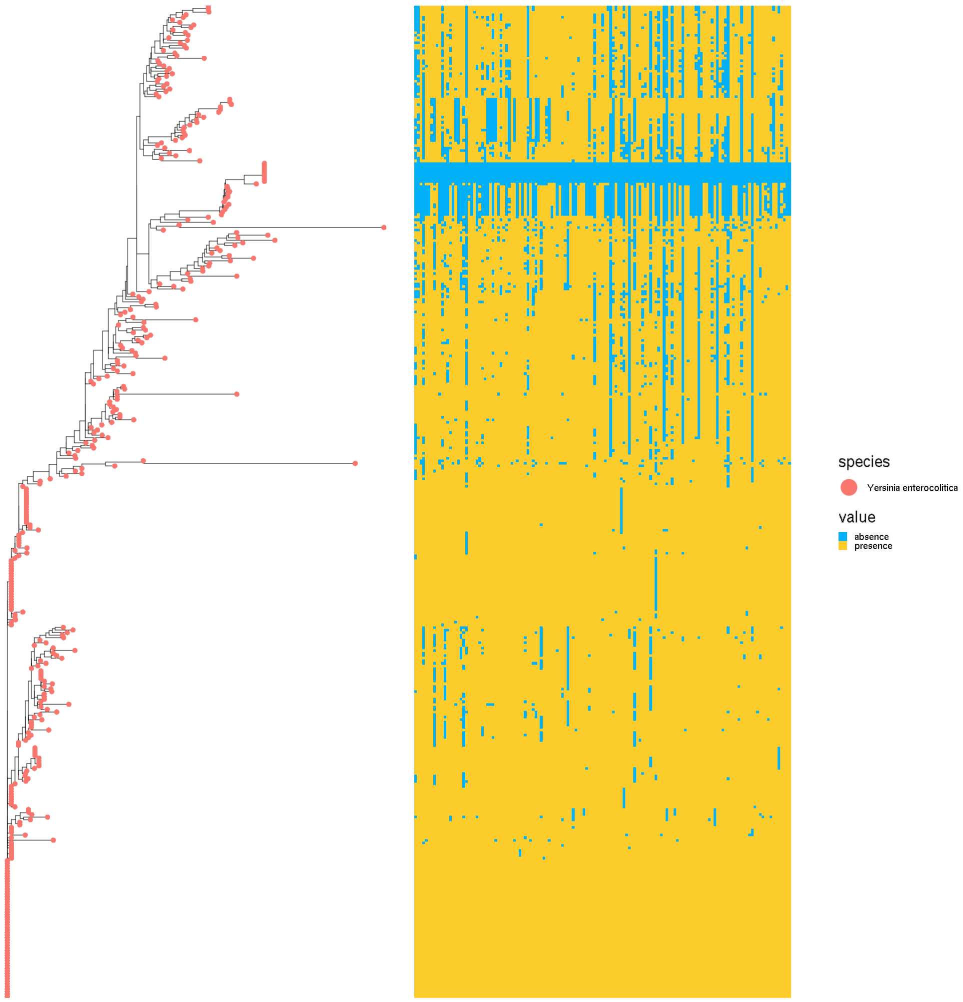


Figure S9: The phylogeny based on the presence/absence of 1000 bp SSR fragments among the entire pan-genome of *Salmonella enterica*. The matrix on the right of the phylogeny shows the SSRs of *Yersinia enterocolitica* of individual strains. Color yellow represents the presence of a region, while blue represents the absence of a region.

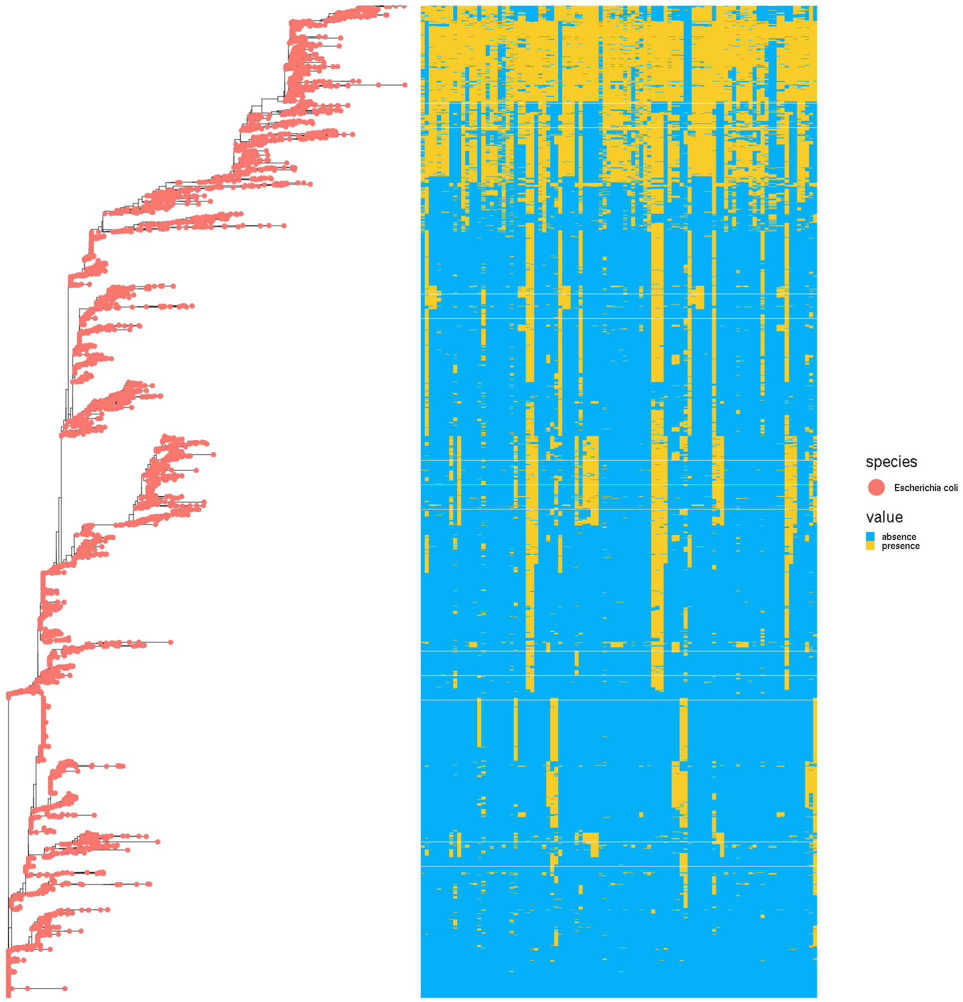


Figure S10: The phylogeny based on the presence/absence of 1000 bp SSR fragments among the entire pan-genome of *Salmonella enterica*. The matrix on the right of the phylogeny shows the SSRs of *Escherichia coli* of individual strains. Color yellow represents the presence of a region, while blue represents the absence of a region.

| Background Strains | Foreground Strains | | | | In silico dataset | | | | | |
|---|---|---|---|---|---|---|---|---|---|---|
| **Background** | **Target Species** | **Target Strain** | **Assembly Accession** | **Accession ID** | **Abundance List** | **0.1 million** | **0.5 million** | **1 million** | **5 million** | **10 million** |
| NC_003212.1 Listeria innocua Clip11262 | Campylobacter_jejuni | RM1221 | GCF_000011865.1 | NC_003912.7 | [0,0.0001,0.001,0.01,0.1] | 5 | 5 | 5 | 5 | 5 |
| NC_004547.2 Pectobacterium atrosepticum SCRI1043 | Clostridium_perfringens | CBA7123 | GCF_002355795.1 | NZ_AP017630.1 | [0,0.0001,0.001,0.01,0.1] | 5 | 5 | 5 | 5 | 5 |
| NC_004557.1 Clostridium tetani E88 | Listeria_monocytogenes | 81-0861 | GCF_000513615.1 | NZ_CP006874.1 | [0,0.0001,0.001,0.01,0.1] | 5 | 5 | 5 | 5 | 5 |
| NC_004668.1 Enterococcus faecalis V583 | Proteus_mirabilis | HI4320 | GCF_000069965.1 | NC_010554.1 | [0,0.0001,0.001,0.01,0.1] | 5 | 5 | 5 | 5 | 5 |
| NC_004917.1 Helicobacter hepaticus ATCC 51449 | Staphylococcus_aureus | RF122 | GCF_000009005.1 | NC_007622.1 | [0,0.0001,0.001,0.01,0.1] | 5 | 5 | 5 | 5 | 5 |
| NC_005139.1 Vibrio vulnificus YJ016 DNA | Vibrio_cholerae | MS6 | GCF_000829215.1 | NZ_AP014524.1 | [0,0.0001,0.001,0.01,0.1] | 5 | 5 | 5 | 5 | 5 |
| NC_006814.3 Lactobacillus acidophilus NCFM | Vibrio_parahaemolyticus | O3:K6 | GCF_000196095.1 | NC_004603.1 | [0,0.0001,0.001,0.01,0.1] | 5 | 5 | 5 | 5 | 5 |
| NC_007575.1 Sulfurimonas denitrificans DSM 1251 | Yersinia_enterocolitica | 8081 | GCF_000009345.1 | NC_008800.1 | [0,0.0001,0.001,0.01,0.1] | 5 | 5 | 5 | 5 | 5 |
| NC_007606.1 Shigella dysenteriae Sd197 | Escherichia coli | 2011C-3493 | GCF_000299455.1 | NC_018658.1 | [0,0.0001,0.001,0.01,0.1] | 5 | 5 | 5 | 5 | 5 |
| NC_008525.1 Pediococcus pentosaceus ATCC 25745 | Salmonella_enterica | USDA-ARS-USMARC-1903 | GCF_000940975.1 | NZ_CP007222.1 | [0,0.0001,0.001,0.01,0.1] | 5 | 5 | 5 | 5 | 5 |
| Note: Above ten strains as the background strains were used for each simulated metagenome. | | | | | | | | | | |

Figure S11: List of foreground and background strains used in generating the *in silico* simulated metagenomic data sets.
